# Blood-based targeted metabolipidomics reveals altered omega fatty acid-derived lipid mediators in relapsing-remitting multiple sclerosis patients

**DOI:** 10.1101/2024.01.04.574253

**Authors:** Insha Zahoor, Jeffrey Waters, Nasar Ata, Indrani Datta, Theresa L. Pedersen, Mirela Cerghet, Laila Poisson, Silva Markovic-Plese, Ramandeep Rattan, Ameer Y. Taha, John W. Newman, Shailendra Giri

**Author notes:** To whom correspondence should be addressed: Shailendra Giri, PhD Senior Scientist Department of Neurology, E&R 4051, Henry Ford Health, 2799 W Grand Blvd, Detroit, Michigan, 48202, USA. Equal contribution (Insha Zahoor, Jeffrey Waters). Equal contribution (John W. Newman, Shailendra Giri).

## Abstract

Unresolved and uncontrolled inflammation is considered a hallmark of pathogenesis in chronic inflammatory diseases like multiple sclerosis (MS), suggesting a defective resolution process. Inflammatory resolution is an active process partially mediated by endogenous metabolites of dietary polyunsaturated fatty acids (PUFA), collectively termed specialized pro-resolving lipid mediators (SPMs). Altered levels of resolution mediators have been reported in several inflammatory diseases and may partly explain impaired inflammatory resolution. Performing LC-MS/MS-based targeted lipid mediator profiling, we observed distinct changes in fatty acid metabolites in serum from 30 relapsing-remitting MS (RRMS) patients relative to 30 matched healthy subjects (HS). Robust linear regression revealed 12 altered lipid mediators after adjusting for confounders (p <0.05). Of these, 15d-PGJ2, PGE3, and LTB5 were increased in MS while PGF2a, 8,9-DiHETrE, 5,6-DiHETrE, 20-HETE, 15-HETE, 12-HETE, 12-HEPE, 14-HDoHE, and DHEA were decreased in MS compared to HS. In addition, 12,13-DiHOME and 12,13-DiHODE were positively correlated with expanded disability status scale values (EDSS). Using Partial Least Squares, we identified several lipid mediators with high VIP scores (VIP > 1: 32% - 52%) of which POEA, PGE3, DHEA, LTB5, and 12-HETE were top predictors for distinguishing between RRMS and HS (AUC =0.75) based on the XGBoost Classifier algorithm. Collectively, these findings suggest an imbalance between inflammation and resolution. Altogether, lipid mediators appear to have potential as diagnostic and prognostic biomarkers for RRMS.

## INTRODUCTION

Multiple sclerosis (MS) is an immune-mediated disease of the central nervous system (CNS), with an incapacitating effect on the functional systems of the patient (Dendrou et al., 2015). The exact trigger behind disease development is still unknown, which complicates its accurate diagnosis, prognosis, and management (Olsson et al., 2017). The present knowledge suggests the role of environmental (non-infectious or infectious), genetic, and epigenetic factors in altering disease risk. A large prospective study has found a significant inverse association between polyunsaturated fatty acid (PUFA) intake and the risk of MS; specifically, the effects were only significant for plant-derived *n*-3 alpha-linolenic acid (ALA) and not for marine *n*-3 fatty acids (Bjornevik et al., 2017; Bjornevik et al., 2019). Thus, based on the outcome of multiple studies, low PUFA intake may be a modifiable risk factor for MS (Parks et al., 2020). This reflects the requirement for in-depth research on dietary factors to understand the definite mechanisms of disease etiopathogenesis.

Unresolved inflammation is a hallmark of MS and several other autoimmune diseases; however, current therapeutic options fail to suppress ongoing inflammation, resulting in inflammatory attacks that gradually increase in severity over time. The underlying disease process is marked by chronic uncontrolled peripheral inflammation and neuroinflammation which often result in demyelination and neurologic deficit in patients due to infiltration of immune cells (Compston & Coles, 2008; Dutta & Trapp, 2011). Thus, targeting mechanisms that can curtail excessive inflammation and promote its timely resolution are of great interest.

Specialized pro-resolving lipid mediators (SPMs) are thought to be critical regulators of chronic inflammation that counterbalance the inflammatory response (Dalli et al., 2013). SPMs are purported endogenous metabolites of n6 and n3 PUFAs, that encompass an array of metabolites including the n6-derived lipoxins and epoxides, and n3-derived epoxides, protectins, maresins and resolvins (Dalli et al., 2013; Fullerton & Gilroy, 2016). While the direct connection between long-chain omega-3 fatty acid intake and SPM production remains unclear (Calder, 2020), the epoxides (epoxyeicosatetraenoic and epoxy docosapentaenoic acids), resolvins, protectins and maresins can be biosynthesized from EPA and docosahexaenoic acid (DHA) through the stereoselective and concerted action of the same enzymes engaged in classical eicosanoid production, namely cyclooxygenase (COX) COX-2, lipoxygenases (LOX) LOX-5, LOX-12, and LOX-15 as well as cytochrome P450 (CYPs)(Dyall et al., 2022; Serhan, 2014) (**Fig. 1**). Altered levels of SPMs have been found in several inflammatory diseases, including periodontitis, asthma, cystic fibrosis, mastitis, atherosclerosis, rheumatoid arthritis, and others (Duvall & Levy, 2016; Elabdeen et al., 2013; Fredman et al., 2016; Zahoor & Giri, 2021; Zahoor et al., 2021).

**Figure 1:**
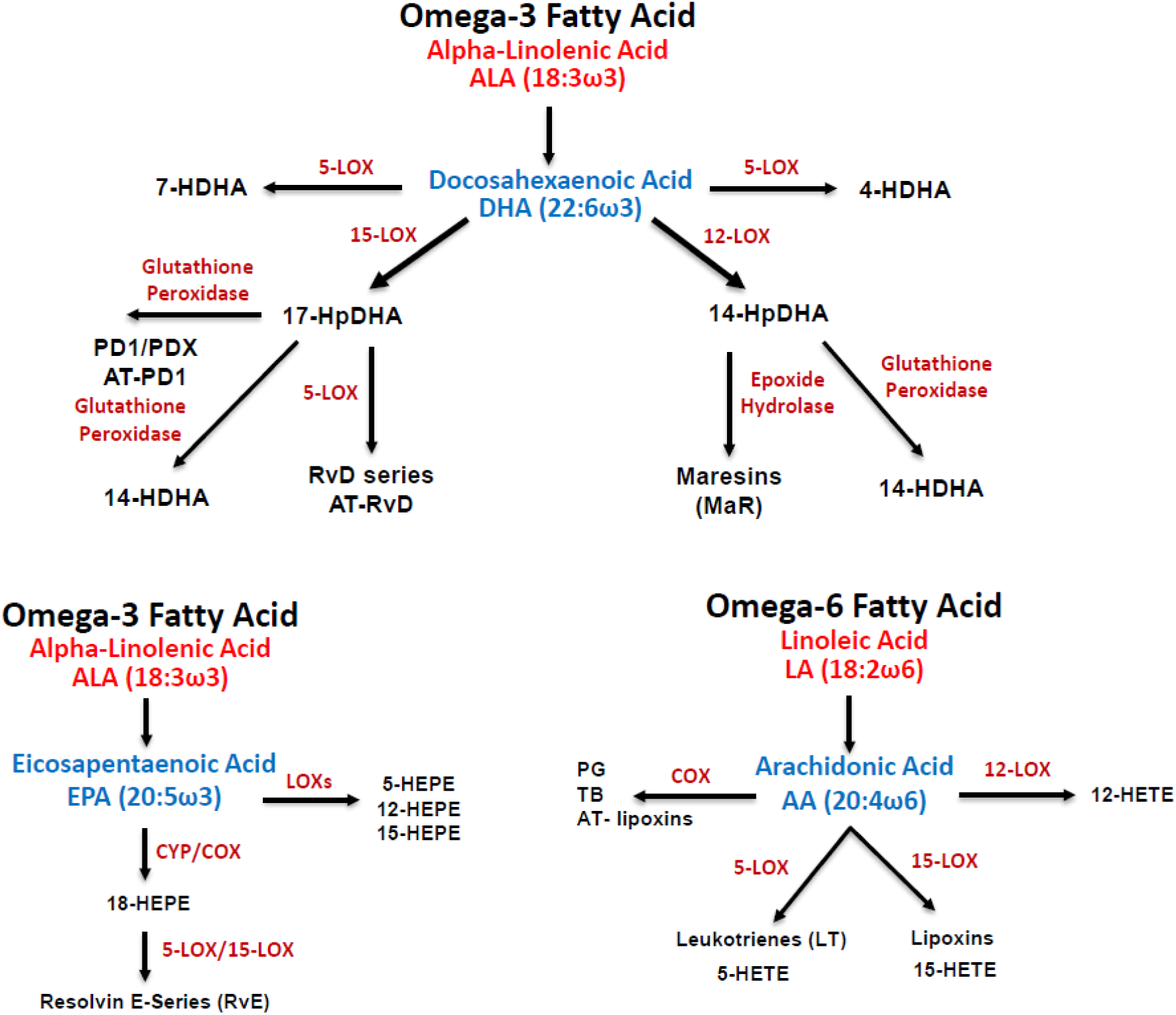
Schematic showing omega fatty acid-derived lipid mediator metabolome, with enzymes (shown in red text), pathway markers, and SPMs. Abbreviations: 5/12/15-LOX, 5/12/15-lipoxygenase; COX, cyclooxygenase; CYP, cytochrome P450

There is currently a debate on whether most SPMS can be reliably measured in open circulation using current analytical techniques (O’Donnell et al., 2023; Schebb et al., 2022), possibly due to their short half-life and rapid elimination (Psychogios et al., 2011; Skarke et al., 2015). However, the immunomodulatory roles for lipoxins and fatty acid epoxides are well established (Chandrasekharan & Sharma-Walia, 2015), and collections of resolvin-, protectin- and maresin-sensitive receptors have been reported (Ferguson et al., 2020). Moreover, the protective ability of SPMs in various models of human diseases has been demonstrated and suggests them as attractive therapeutic candidates in MS, as revealed by metabolomics approaches (Zahoor & Giri, 2021; Zahoor et al., 2021). Moreover, changes in oxylipin precursors to these metabolites can be measured in plasma and are thought indicative of the activation of these pathways (Marchand et al., 2023; Mozurkewich et al., 2016). Specifically, 18-hydroxyeicosapentaenoic acids (18-HEPE), 17-hydroxydocosahexaenoic acid (17-HDHA), and 14-HDHA have been reported as markers for the activation of the E-series resolvins, D-series resolvins, and maresin biosynthetic pathways, respectively (Abdulnour et al., 2014; Barden et al., 2014; Weylandt et al., 2012).

Omega-3 metabolites, specifically, DHA derived SPMs have been reported to be low in the serum of MS patients (Kooij et al., 2020; Pruss et al., 2013) and experimental autoimmune encephalomyelitis (EAE) mouse models (Derada Troletti et al., 2021; Mangalam et al., 2013; Poisson et al., 2015; Sanchez-Fernandez et al., 2022; Zahoor et al., 2023). Indeed, some clinical trials suggest that omega-3 supplementation may be beneficial for MS patients (Ramirez-Ramirez et al., 2013; Torkildsen et al., 2012; Weinstock-Guttman et al., 2005). On the other hand, restoring the level of long-chain omega-3 fatty acids by supplementation in relapsing-remitting MS (RRMS) patients (Torkildsen et al., 2012) showed no improvement in neurological symptoms (Farinotti et al., 2012; Parks et al., 2020; Ramirez-Ramirez et al., 2013; Riccio et al., 2016; Torkildsen et al., 2012; Weinstock-Guttman et al., 2005), raising the question as to why omega-3 supplementation is not effective in MS? It could be postulated that in MS patients, there may be defects in the omega-3 fatty acid (specifically DHA) metabolism that result in a deficiency of downstream SPM metabolites, leading to unresolved chronic inflammation and a delay in the healing/repair process, thus resulting in continued neuronal damage. However, limited information is available regarding the presence or abundance of SPMs (e.g., epoxides, resolvins, protectins, and maresins) in MS patients (Zahoor & Giri, 2021). Thus, we hypothesize that SPMs and their associated pathway markers play an important role in MS, and identification of specific DHA-derived metabolite markers of SPMs may lead to a better manipulation of the omega-3 fatty acid metabolism for treatment of MS patients. With this aim, we profiled the PUFA-derived lipid mediators including SPMs in serum samples obtained from RRMS patients and matched healthy controls (HC) using targeted lipidomics to determine if there is a failure to produce SPMs due to impaired omega-3 PUFA metabolism in RRMS patients. Our present study provides additional evidence in support of the importance of omega-3 PUFAs and identifies an imbalance between the levels of pro-resolution and pro-inflammatory lipid mediator in MS patients.

## MATERIALS AND METHODS

### Patient recruitment and sample collection

A total of 30 blood samples collected from unrelated relapsing-remitting (RRMS) patients were used for the present study from the Biorepository at the University of California San Francisco (UCSF). Patients were diagnosed with MS, according to the 2001 International Panel Diagnostic Criteria (McDonald et al., 2001). The study stands approved by the Institutional Review Board of UCSF (#10-00104) and Henry Ford Hospital (#12829). All the patients/guardians were informed about the study and written informed consent was acquired from them by the ethical standards laid down by the World Health Organization (WHO) and Declaration of Helsinki 1964 and its later amendments or comparable ethical standards. Patients were treatment-naive at the time of sample collection. Inclusion criteria for selecting patients included clinico-radiological confirmed MS patients. In contrast, exclusion criteria included patients who did not satisfy diagnostic criteria, patients with previous history or family history of other neurodegenerative and/or inflammatory diseases, and patients who rebutted from participating in this study. Simultaneously, serum samples from 30 age and sex-matched healthy controls (HS) (to confine the confounding effect) were also used to serve as controls. All subjects including, RRMS patients and HS, were residents of the United States. Inclusion criteria for the selection of controls included healthy individuals with minor neurological problems like back pain and headache and those with no previous or family history of MS or any other autoimmune disease.

### Serum collection

For serum analysis, blood samples from RRMS patients and HS were collected in red-capped vacutainer tubes. The serum was separated from blood by centrifugation at 1500 rpm for 10 min. The clear yellow liquid supernatant was collected from the top and stored in respective yellow-capped serum tubes at -80°C till further processing. The samples were shipped in dry ice to the site of the study.

### Liquid chromatography-mass spectrometry (LC-MS/MS)-based lipid mediator and anti-inflammatory drug profiling

Serum lipid mediators, endocannabinoids, unesterified PUFAs, ibuprofen, and acetaminophen were quantified after protein precipitation in the presence of isotopically labeled analytical surrogates by LC-MS/MS as recently described (Pedersen et al., 2021). Briefly, human serum samples were thawed in a rack over wet ice, 50 µL of serum were added to a 96-deep well polypropylene plate containing 5 µL of 10,000nM labeled oxylipin class surrogates, 5 µL of 5 µM 1-cyclohexyl ureido, 3-dodecanoic acid (CUDA) and 1-phenyl 3-hexadecanoic acid urea (PHAU) internal standard, and 5 µL of 0.2 mg/mL butylated toluene (BHT)/ethylenediaminetetraacetic acid (EDTA) antioxidants, which was followed by swirling to mix (Pedersen et al., 2021). A 185 µL volume of 50:50 methanol:acetonitrile was then added for a total volume of 250µL, capped with a non-slit silicon mat, and vortexed for 30 sec. The plate was placed in a -20 °C for 30 min and insolubles and proteins were pelleted by centrifugation for 10 min at 2000 g at 4 °C. It was followed by placing a filter plate on top of a conical well 450 µL microtiter plate, set in wet ice. Using a multi-channel pipettor, 100-150 µL of sample supernatant was put into a filter plate, paying attention to avoid particulate. The filter plate was centrifuged for 3 min at 500 g and 4 °C, checking that the wells eluted completely. The filter plate was removed, and the microtiter plate was capped with a thermally sealed (polypropylene-backed) foil cover. The plate was transferred to the instrument for acquisition or stored at -20 °C, wrapped in foil for up to 7 days. Oxylipins including SPMs, endocannabinoids, unesterified PUFAs, and non-steroidal anti-inflammatory (NSAIDs) medications were analyzed by electrospray ionization mode (ESI) mode on a Sciex 6500 QTrap coupled to a Shimadzu 30AD UPLC system as previously detailed (Pedersen et al., 2021). Relative concentrations were obtained for a subset of endocannabinoids and all unesterified PUFAs (linoleic acid, alpha-linolenic acid, arachidonic acid, eicosapentaenoic acid, docosahexaenoic acid) and absolute concentrations (nM) were obtained for all other targets.

### Statistical analysis and machine learning Quality control

Metabolite concentrations were obtained from external standard curves and using surrogate standards to correct for analyte recovery. All statistical analyses were performed using the Python programming language (Chong et al., 2019; Seabold, 2010; Van Rossum, 2009). Metabolites with missing values of > 30% were dropped from the study, while the remaining missing values were inputted using the k-nearest neighbors algorithm (n =3) (Altman, 1992).

Metabolite concentrations were log-transformed and for heatmaps and dimensionality reduction, Z-scores were calculated from the log-transformed values.

### Robust linear regression and correlation analysis of lipidomic profiles between RRMS patients and HS

Per metabolite, comparisons were made between RRMS and HS via robust linear regression using the HuberT method to control for noise, while controlling for age, sex, and drug treatment (Ibuprofen/acetaminophen). Significant differences based on robust linear regression were determined at p <0.05 for the coefficient corresponding to the difference between HS and RRMS and corrections for multiple comparisons were calculated using the Holm-Sidak method. Both corrections for multiple comparisons and linear regression were carried out using the statsmodels Python packages (Seabold, 2010). Correlation with EDSS scores was performed on RRMS samples using the biweight mid-correlation while controlling for age, sex, and treatment (Vallat, 2018). For visualizations, heatmaps for both all metabolites and those that were significant based on robust linear regression were generated using Z-scores of the log-transformed values. Visualizations for significantly correlated lipids (p <0.05) were done via robust linear regression plots. In addition, a heatmap displaying correlation coefficient values for all lipid mediators that were significant for EDSS or one of the covariates (Age, Sex, ibuprofen, acetaminophen) was plotted. All plots were made using the Seaborn and Matplotlib Python libraries (Hunter, 2007; Waskom, 2021). All data wrangling was done using numpy and pandas (Harris et al., 2020; Reback, 2021).

### Dimensionality reduction analysis of metabolites

To identify overarching patterns in the data set, as well as key lipids associated with these patterns, principal component analysis (PCA) and partial least squares discriminant/regression analysis (PLSDA/PLSR) were used via the sci-kit learn library. For PLSDA, the target feature used was group (HS vs RRMS) while for PLSR only RRMS samples were used with the target feature being EDSS. R2/Q2 values in conjunction with leaving one out cross-validation were used to determine the prediction strength of the components. Loading score plots were generated for both PLSDA and PLSR with VIP scores, parent fatty acid (PFA), and the enzyme that catalyzes the reaction between lipid and its parent fatty acid being overlayed via color labeling. All dimensionality reduction was done using the sci-kit learn library ^44^.

### Clustering and correlation network analysis of lipid mediators

The clustering of lipid mediators was performed using hierarchal clustering. First, a correlation matrix was generated using the biweight mid-correlation method, next the matrix was plotted as a heatmap with columns and rows being sorted by hierarchical clustering using a Euclidean distance metric and ward method for linkage. The optimal cutoff for clusters was determined visually from the heatmap. To determine which lipids had similar intensity patterns, correlation networks were generated based on a biweight mid-correlation matrix of all lipids evaluated in the study. For the network, edges exist between lipids if the p-value <0.05 and the correlation coefficient was ≥0.4, isolated nodes were eliminated. Lipids were color-labeled by PFA and by the enzyme that catalyzes the reaction between the lipid and its PFA. For determining altered expression patterns between lipids derived from the same PFA or enzyme, interconnectivity scores were calculated and compared between groups (HS vs RRMS) by lipids in the correlation network with significance determined via a permutation test; here interconnectivity was defined as the ratio of edges between members of the same groups (enzyme/PFA) within the correlation network and the maximum number of connections possible. Clustering and correlation networks were done using Scipy, sci-kit Learn, and network libraries (Pedregosa, 2011; Virtanen et al., 2020).

### Multivariate analysis for classification and regression

To see if EDSS and classification (HS or RRMS) could be determined based on lipid expressions, the XGBoost algorithm was used. First lipids with VIP scores >0.8 were identified from PLSDA. Top lipids were then selected using XGBoost and leave one out cross validation. Model hyperparameters were then optimized using the grid search function from the sci-kit learn library (Pedregosa, 2011). For the final classification model, regularization via the reg_lambda parameter was implemented (reg_lambda=1) to mitigate the effects of noise and to improve generalization. After determining the top lipids and optimal hyperparameters a final model was generated, and performance was evaluated using leave-one-out cross-validation and the area under the curve (AUC) metric. A similar pipeline was used for regression with EDSS (using PLSR VIP scores rather than PLSDA). For visualizing model predictions, a receiver operating characteristic (ROC) curve was generated. Furthermore, the prediction space for the final model was translated and plotted into the low dimensional space generated from PLSDA (where the space generated by PLSDA was based on the expression of only the top 5 lipids) and loading scores for the top lipids were plotted as well as a heat map showing the expression patterns of top lipids across RRMS and HS samples using Z-scores. Analysis was done using the XGBoost and sci-kit learn libraries (Pedregosa, 2011) and plotting using matplotlib and seaborn (Hunter, 2007; Waskom, 2021).

## RESULTS

### Baseline characteristics of the study population

To identify whether differential expression patterns exist in lipid mediators between RRMS and HS, we analyzed 90 lipid mediators from serum samples of 30 RRMS and 30 HS. On average, participants were aged 34.3 ± 8.35 years (**Table 1**) and were mostly female (73.3%) of the Caucasian race (71.7%). RRMS had disease durations averaging 1.43 years were not treated with any disease-modifying therapies (DMTs) and were moderately disabled (median EDSS score =2.0).

**Table 1.** Demographic and clinical data of study cohorts.

| **Variable** | **HS** | **RRMS** |
| --- | --- | --- |
| **No. of subjects** | N=30 | N=30 |
| **Mean age ± SEM** | 27.87±5.23 | 34.3±8.35 |
| **Female/Male** | 22/8 | 22/8 |
| **Longest Disease Duration** | N/A | 8 |
| **Mean EDSS** | N/A | 2.27 |
| **Race** | Afm(2)/<br>white_hispanic(3)/<br>white_non_hispanic(25) | Afm(4)/<br>white_hispanic(8)/<br>white_non_hispanic(18) |

### Lipid mediator alterations between RRMS and HS and association with disease severity

Visually, overall expression of lipid mediators in serum suggests considerable differences between RRMS and HS (**Fig. 2A**). Robust linear regression (after controlling for age, sex, and ibuprofen/acetaminophen levels) showed 12 significantly altered lipid mediators (p <0.05), of which 3 were increased, including 15-deoxy-delta12,14-prostaglandin J2 (15d-PGJ2), leukotriene B5 (LTB5), prostaglandin E3 (PGE3) and 9 were decreased, including Prostaglandin F2alpha (PGF2a), 8,9-dihydroxyeicosatrienoic acid (8,9-DiHETrE), 5,6-dihydroxyeicosatrienoic acid (5,6-DiHETrE), 20-hydroxyeicosatetraenoic acid (20-HETE), 15-hydroxyeicosatetraenoic acid (15-HETE), 12-hydroxyeicosatetraenoic acid (12-HETE), 12-hydroxyeicosapentaenoic acids (12-HEPE), 4-hydroxydocosahexaenoic acid (14-HDoHE) and docosahexaenoyl ethanolamide (DHEA) in RRMS samples compared to HS (**Fig. 2B-C**). Of these lipids, the 12/15-LOX-derived 12-HETE, 12-HEPE, and 14-HDoHE showed the greatest difference between the control and case with adjusted p-values <0.05. Associations with disease severity (EDSS) and lipids were examined using the biweight mid-correlation while controlling for the patient’s age, sex, and ibuprofen/acetaminophen levels. The linoleate-derived 12,13-dihydroxy-9-octadecenoic acid (12,13-DiHOME) and alpha-linoleic acid-derived 12,13-dihydroxyoctadeca-9,15-dienoic acid (12-13-DiHODE) were found to be positively correlated with EDSS (p <0.05) (**Fig. 3A-B**) with adjusted p-values 0.94 and 0.69 for 12,13-DiHOME and 12,13-DiHODE respectively while several COX and LOX products negatively correlated (p <0.05) with ibuprofen levels (**Fig. 3C**). In general, ibuprofen and acetaminophen levels were evenly distributed across both HS and RRMS samples **(Supp. Fig. 2)**.

**Figure 2:**
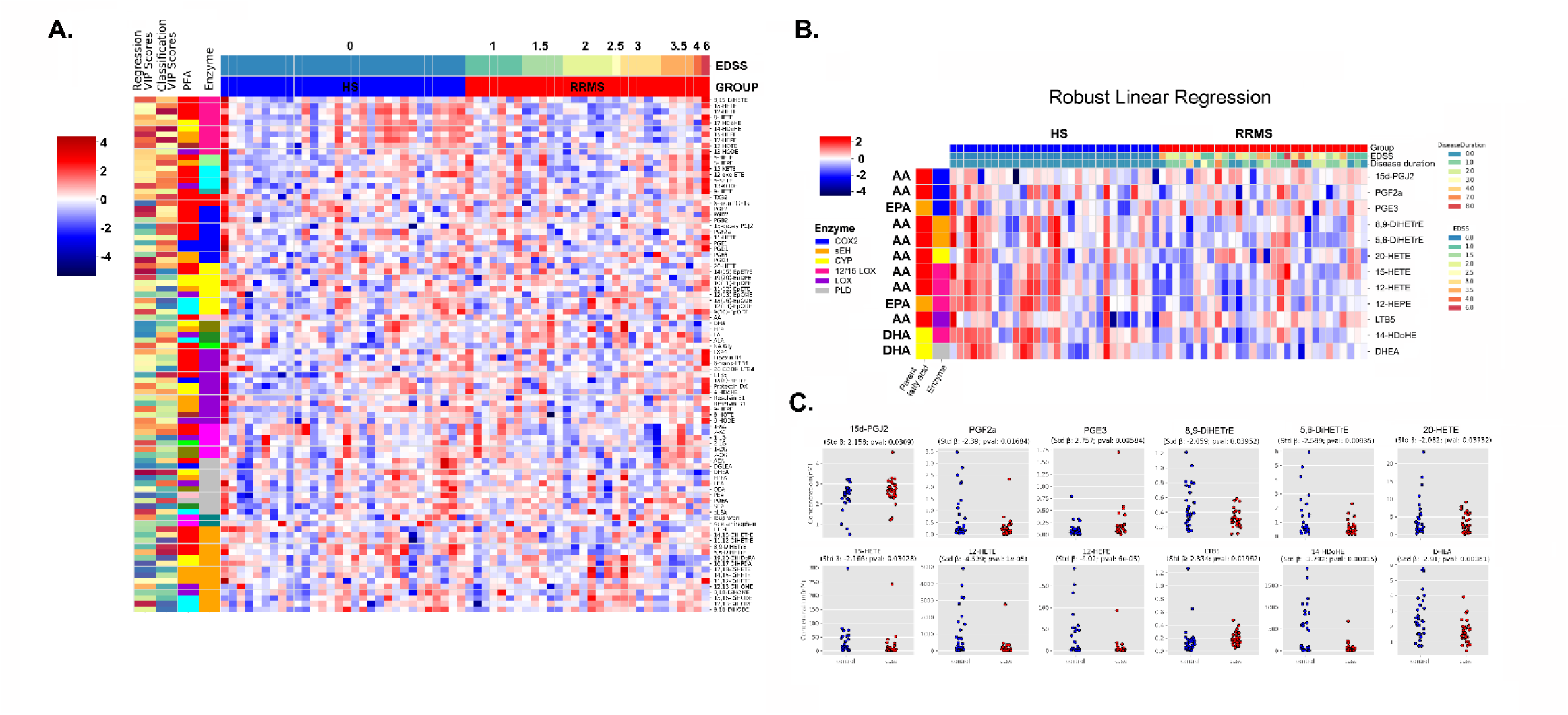
Heatmap showing z-scores of log concentration of lipids. **A.** Lipids are ordered first by the enzyme that catalyses the reaction between the lipid and its parent fatty acid, then by its parent fatty acid, followed by its VIP score from PLS-DA and PLSR, where the Z score varies from 4 to -4. Healthy control on 0 and the RRMS from 1 to 4.6 **B.** The heatmap of z-scores derived from the log expression values of lipids which were deferentially expressed between HS and RRMS based on the coefficients from linear regression with M-estimation. **C.** Scatter plots depicting the concentration (nM) of significantly altered lipids between HS and RRMS (n=30 each).

**Figure 3:**
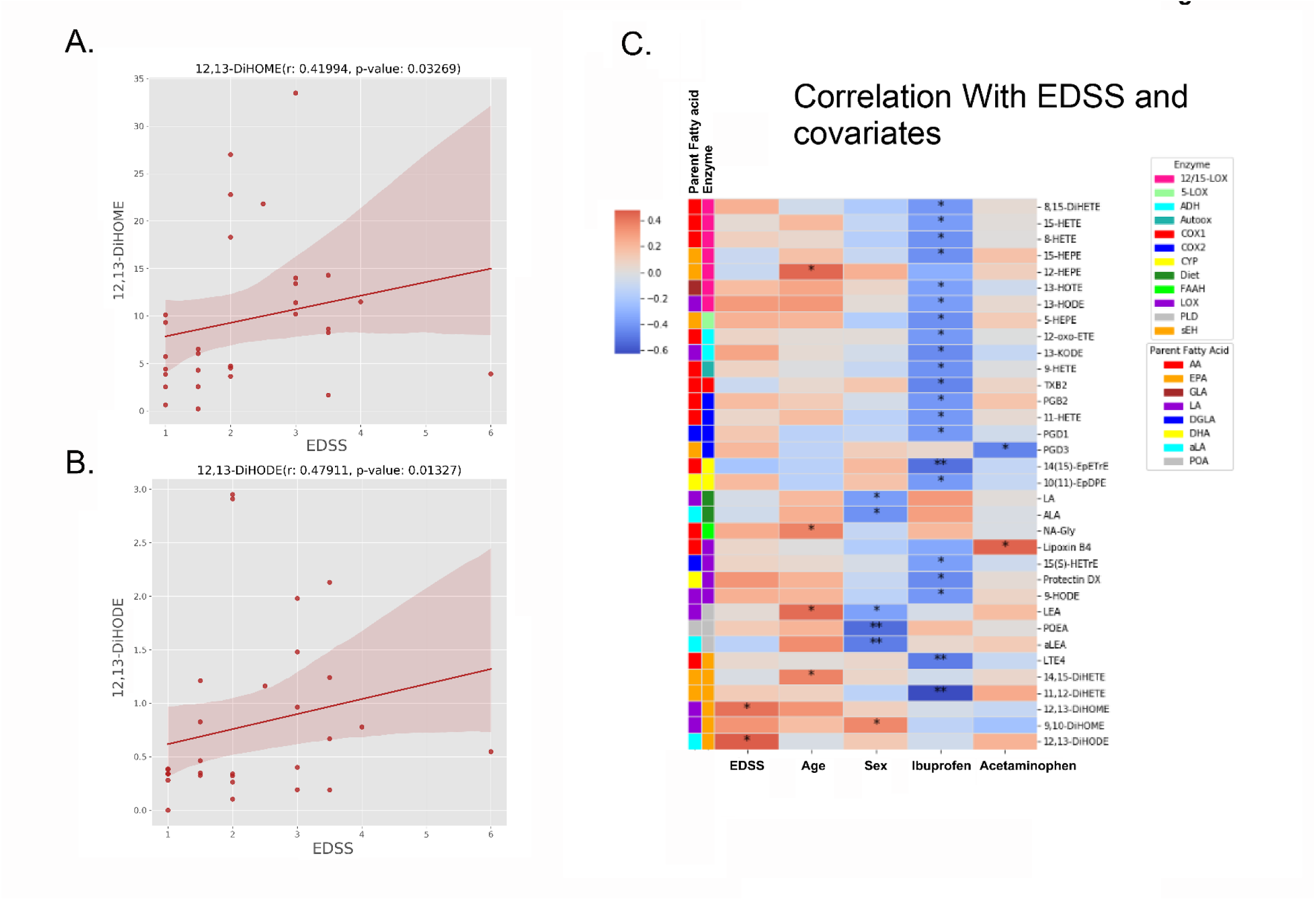
12,13-dihydroxy-9-octadecenoic acid (12,13-DiHOME) and (9Z,15Z)-12,13-dihydroxyoctadeca-9,15-dienoic acid (12-13-DiHODE) showed a positive correlation with severity of the disease. **A-B.** Scatter plots show the association of RRMS disability scores (EDSS) on the X-axis and concentration (nM) of 12,13-DiHOME or 12-13-DiHODE on the Y-axis. Each dot represents a single RRMS patient. In addition, the line of best fit generated from the linear regression model using M-estimation is plotted as well as the 95% CI bands. The correlation coefficient(r) and the p-value from the correlation between the lipid and EDSS while controlling for age and sex using the biweight mid-correlation test are listed above the figure. **C.** The z-scores for the log concentrations (nM) for all lipids were significantly correlated (p-value < 0.05) with at least one of the variables including EDSS, age, sex, ibuprofen, and acetaminophen.

### Dimensionality reduction analysis of lipid mediators

To observe if overarching patterns exist between RRMS and HS, initially, we performed PCA to determine whether patterns could be observed with an unsupervised approach. We observed poor separation of RRMS from HS based on 3 components with a total explained variance of 49.1% (**Supp. Fig. 1)**. However, an improved separation was observed when using the first 3 components of PLSDA, suggesting an underlying covariance structure associated with sample classification (**Fig. 4A**). Visually, loading scores for PLSDA showed some degree of clustering based on enzyme (**Fig. 4B**). Interestingly, 14-HDoHE, 12-HETE, and 12-HEPE showed a strong association with lower levels in RRMS compared to control as well as having the highest VIP scores for PLSDA (2.28, 2.25, 2.19 respectively) with total VIP scores >0.8 =32%. Furthermore, all 3 lipids are metabolized by 12/15 LOX and show similar expression patterns across most samples along with other lipids evaluated in this study that are also metabolized by 12/15 LOX, despite being derived from different PUFAs (**Fig. 1A**). PLSR of RRMS samples using EDSS as the target variable revealed spatial patterns within the low dimensional space constructed by the first 3 components of PLSR which seemed to correspond to the progression of disease severity with a sizeable degree of variance in EDSS scores from 1-3 (**Fig. 4C**). Notably, scores 3.5 and 4, visually, show distinct clustering with each other, suggesting that these scores may correspond to a more uniform point in disease severity concerning the lipid mediator expressions.

**Figure 4:**
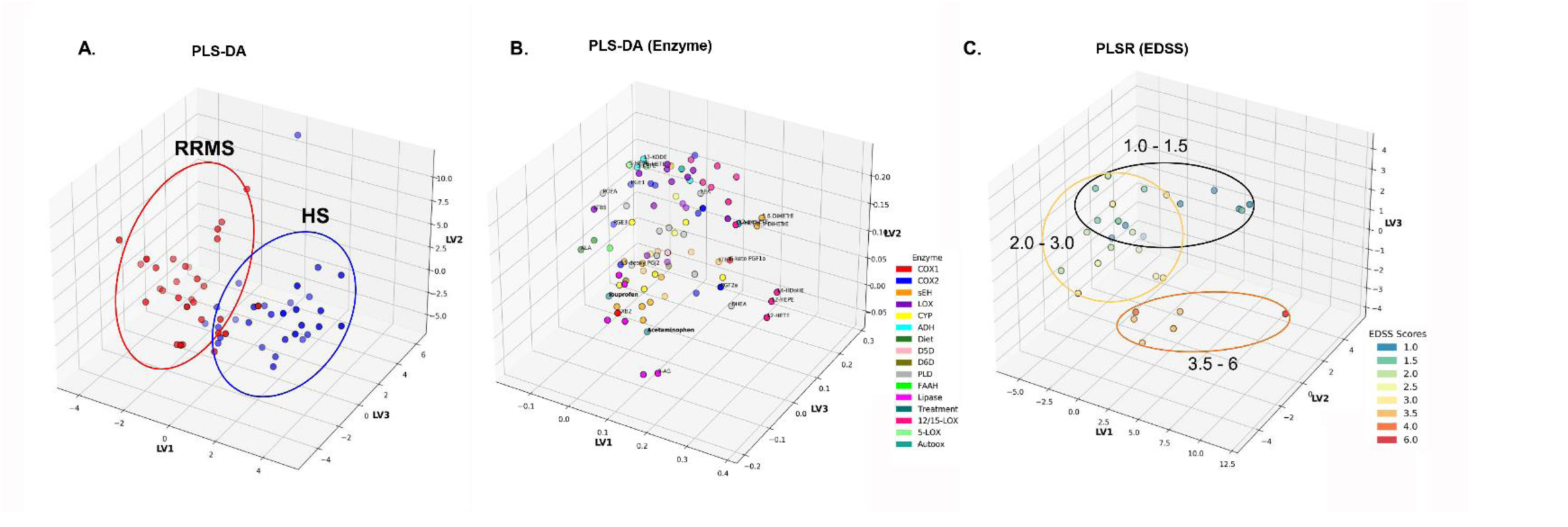
PLSR plots showing lipids and enzyme-based catalytic reaction proving the different levels of RRMS level. **A.** Multivariate analysis of significantly altered lipids using 3D score plot of PLS-DA for HS and RRMS groups. **B.** Loading scores for PLS-DA, having Lv1 vs Lv2 vs Lv3 for enzymes, in which each lipid is grouped by color based on the enzyme that catalyzes the reaction from the parent fatty acid. Individual lipids with VIP (variable importance in projection)> were used for analysis. **C.** Loading scatter plot for the partial least-squares regression (PLSR) of RRMS samples on disease disability score (EDSS). RRMS patients were color-labeled based on EDSS scoring.

### Clustering analysis, correlation networks, and classification of samples based on lipid mediator expression

To see if a highly predictive classification model could be constructed based on the lipid mediator profile, we used the XGBoost algorithm (T;). We chose this classification model due to its robustness to outliers, ability to effectively handle high-dimensional metabolomic data, and its feature importance function which can aid in identifying key metabolites that differentiate between MS patients and healthy subjects, providing invaluable insights for biological interpretation. Classification of lipid mediators using XGBoost which stands for extreme gradient boosting, is a scalable, distributed gradient-boosted decision tree (GBDT) machine learning library. It also has a few hyperparameters that can be tuned to improve model performance, including the learning rate, depth of the trees, and regularization parameters, revealed 5 highly predictive metabolites (AUC =0.75) (**Fig. 5A**) including LTB5, 12-HETE, POEA, DHEA, and PGE3. PLSDA of samples using these 5 lipids generated a very similar sample layout showing regression plotted for lipids that were significantly correlated with EDSS while controlling for covariates (Age, Sex, Ibuprofen, and Acetaminophen with r =0.419 and p =0.032 (**Fig. 5B-C**) compared to the one using all lipid mediators, further suggesting that these 5 lipids are strong representatives of the differences in the lipid mediator profiles between HS and RRMS. Of these 5 lipid mediators, LTB5, POEA, and PGE3 were elevated in the majority of RRMS subjects, while 12-HETE and DHEA were decreased (**Fig. 5D**). Regression with XGBoost performed poorly (average rose =0.43, std dev =0.62), we suspect because of the large variation present in EDSS scores 1-3.

**Figure 5:**
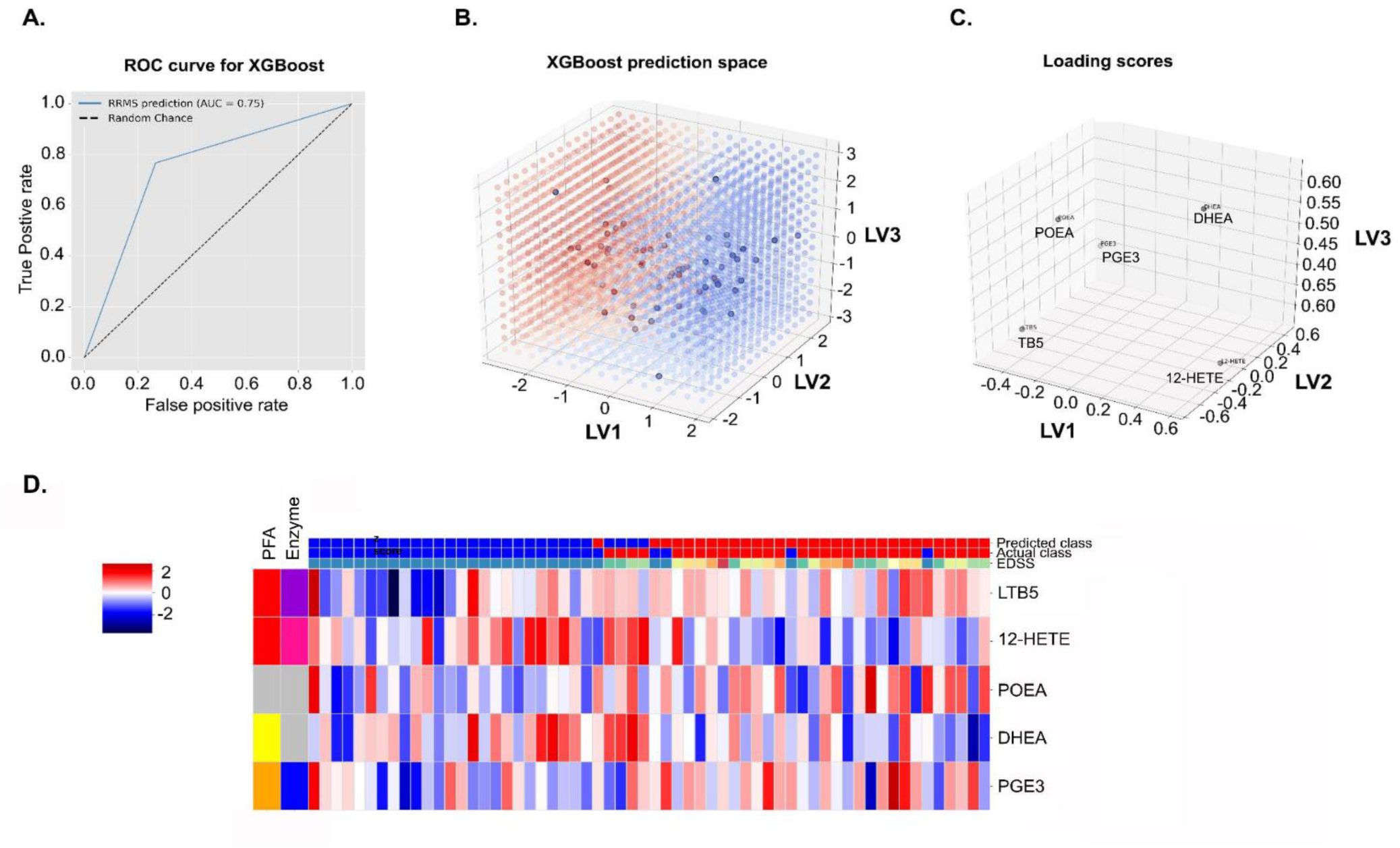
Prediction of true or false positive classification of MS vs. HS using XG Boost. **A.** ROC curve of final XGBoost model with AUC of 0.75 showing higher random positive value than random negative concerning true positive and false positive levels. **B.** XGBoost prediction space projected into the first 3 components of PLS-DA, where the blue color concentration is for the HS and the Red color specifies RRMS having a value ranging from 1.5-3 EDSS which can be a bit low. **C.** Loading scores of most predictive lipid mediators selected by XGBoost for Lv1, Lv 2, and Lv3 show correlations between the indicator and the estimated latent variable score. **D.** The heat map of Z-scores of log expression values of the top lipid mediators used by XGBoost of HS and RRMS.

To evaluate which set of lipids were similar in profile to the 5 highly predictive metabolites identified by XGBoost, hierarchal clustering was performed, and 8 distinct clusters were identified **(Fig. 6A, 6B)**. Most notably, DHEA and POEA were highly associated with fatty acid ethanolamides, while 12-HETE was associated with other 12/15-LOX metabolites. Further exploring lipid mediator associations with each other, we investigated correlation patterns between all lipids in the data set and how these correlations change between HS and RRMS **(Fig. 7A, 7B)**. In general, correlations between lipids were strongest among lipids metabolized by the same enzyme. However, interesting discrepancies were observed among lipids metabolized by PLD as we detected a statistically significant drop in interconnectivity (p =0.01) between lipids metabolized by PLD from HS and RRMS (**Fig 7C**). This finding is corroborated by expression patterns in these lipids as the expression is more uniform in HS – despite each lipid mediator being derived from a different PFA - compared to RRMS. Furthermore, when evaluating lipid interconnectivity by PUFA, we observed a convergence of interconnectivity that occurred in RRMS from HS (**Fig 7D**).

**Figure 6:**
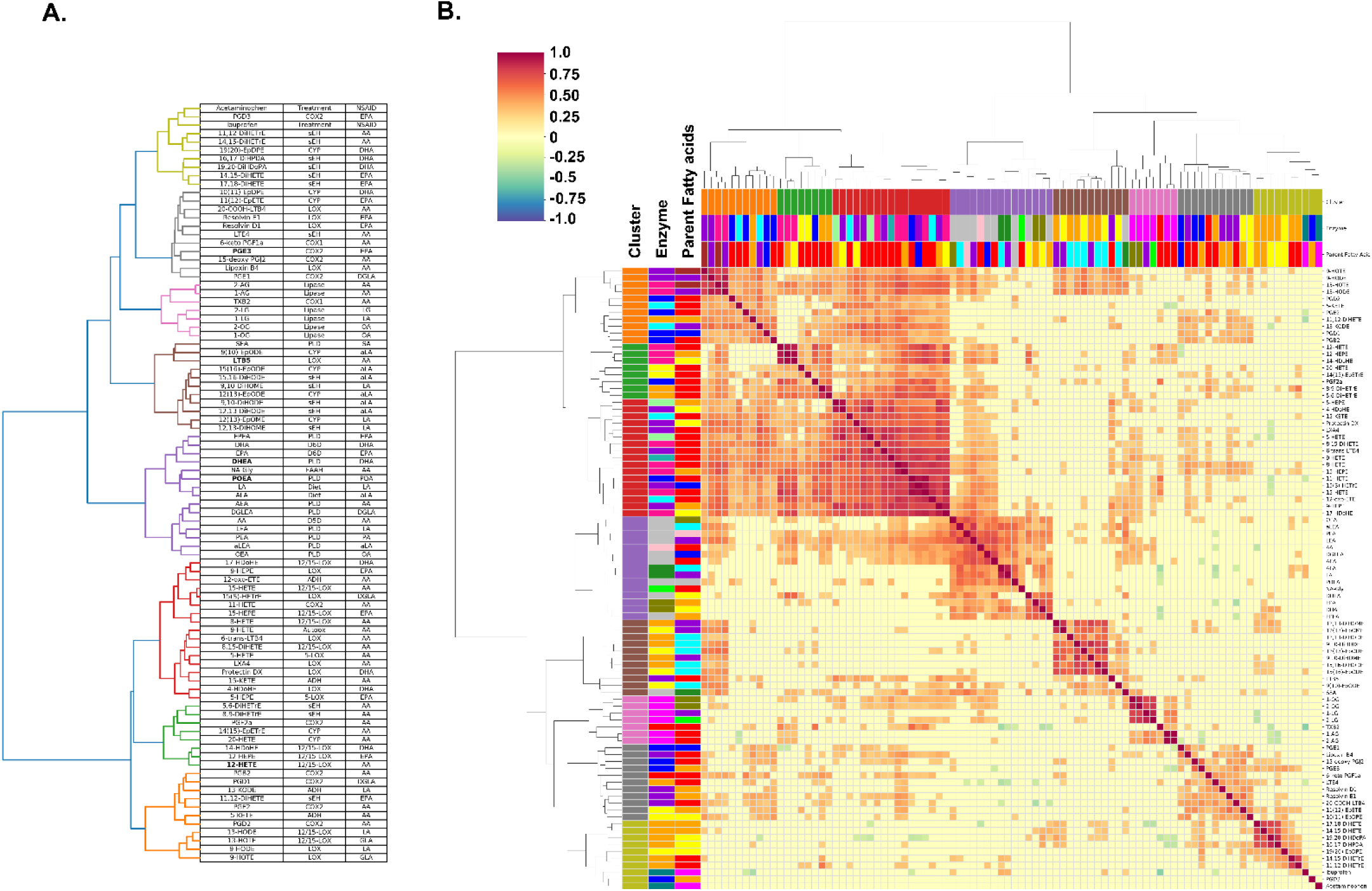
Clustering of lipid mediators based on hierarchical clustering of the biweight mid-correlation matrix in HS and RRMS. **A.** Hierarchical clustering presented in dendrogram table. **B.** Hierarchical clustering presented as a heatmap where color values correspond to the value of the correlation coefficient from the biweight mid-correlation, the value or r ranging from 1 to -1 showing cluster, enzyme, and fatty acid. For hierarchical clustering, the distance metric used was Euclidean and linkage was determined using the average method. Clusters of Green, Red, and Orange are mostly LOX-derived lipids. Purple contains all lipids which are metabolized by PLD. Pink contains all lipids which are metabolized by lipases. Brown and Olive contain lipids metabolized by sEH and CYP but are separated based on the parent fatty acid they are derived from. Brown is mostly derived from alpha-linolenic acid and Olive is EPA and DHA and contains both drugs.

**Figure 7:**
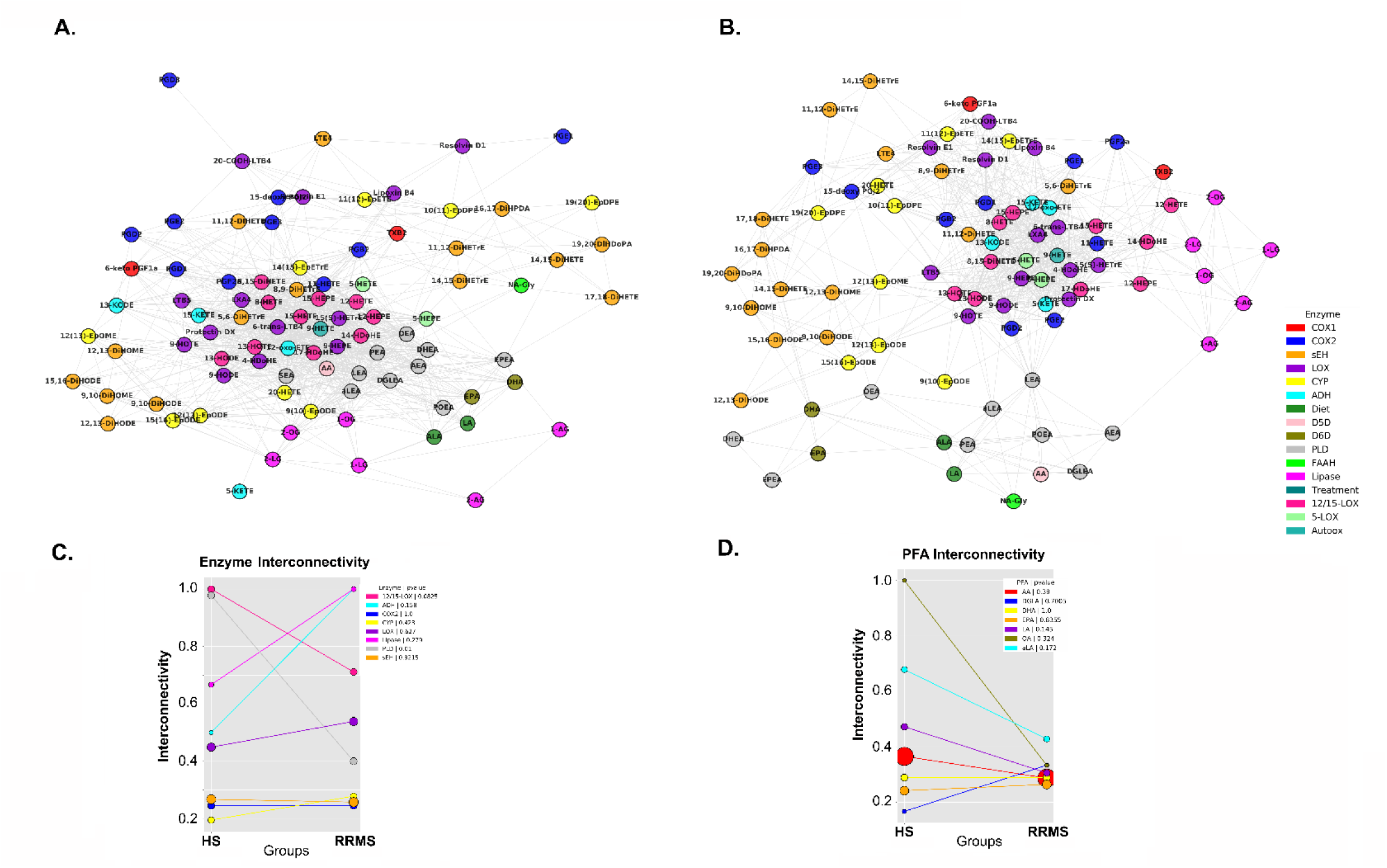
correlation networks for lipids in HS and RRMS. Correlation analysis showing lipid-based network and drug influence across subjects showing a loss in interconnectivity between HS (**A**) and RRMS (**B**). Lipids are color-labeled based on the enzyme which catalyzes the reaction between the lipid and its parent fatty acid. Node proximity is based on edge weight so that groups of nodes that are highly correlated with each other have closer proximity to each other. **C-D.** Interconnectivity values for each lipid group based on enzyme and parent fatty acid, respectively.

## Discussion

The prognosis of MS remains highly unpredictable and is largely determined by the clinical phenotype exhibited by each patient. Thus, an active area of investigation is to identify novel biomarkers that can advance current therapeutic options to treat MS or reverse the disease course. The clinical outcome of MS patients may not be determined solely by the levels of pro-inflammatory mediators but also by mediators of inflammatory resolution. We postulate that endogenous mechanisms to resolve inflammation are intact but become defective in patients with RRMS, leading to sustained inflammation and increased disease progression and disability over time. SPMs can counterbalance the inflammatory response and reduce associated tissue damage in various disease models (Dalli et al., 2013). Therefore, SPMs as critical regulators of chronic inflammation and its associated disability, play a vital role in resolving the disease and are attractive therapeutic candidates for MS. Importantly, if defects in SPM biosynthesis are present, SPM-based therapeutic approaches may be more effective than PUFA supplementation.

In the present study, we provided a broad characterization of the omega-3 fatty acid-derived oxylipin profile alongside other PUFA metabolites, in serum samples of MS patients. Our study reports that the lipid mediator profile in serum was different between HS and RRMS patients, but the detectable SPMs (namely lipoxins A4 and B4, resolvins E1 and D1, and Protectin Dx) did not differ among the groups. However, RRMS patients had increased 15d-PGJ2, PGE3, 11(12)-EpETrE, ALA, 5-HEPE, and LTB5 and decreased DHEA 8,9 DiHETrE, 5,6 DiHETrE, 12-HETE, 12-HEPE and 14-HDoHE levels (**Fig. 2**). interestingly, the 12-LOX metabolites (i.e.,12-HETE, 12-HEPE, and 14-HDoHE) were consistently suppressed in the serum of MS patients. The 12-LOX is activated during platelet degranulation and stimulation of blood clotting, leading to a significant serum enrichment in these metabolites relative to plasma (Pedersen et al., 2021). Thus, reduced levels of these mediators in RRMS relative to HS point to decreased 12-LOX activation in MS patients.

It’s worthwhile to mention here that the levels of these metabolites can be influenced by clotting in serum, thromboxane levels do not cluster with these metabolites. Moreover, in a metabolomics twin study of lipid mediators in serum the variance in the 12-LOX metabolites was the most heritable, with genetics explaining 49% to 73% of their variance (Bermingham et al., 2021). Since platelet 12-LOX-dependent 14-HDoHE formation has been reportedly linked to MaR1 production by neutrophils (Abdulnour et al., 2014), it is interesting to speculate that a reduced 12-LOX activity could lead to a reduced capacity of pro-resolving maresin biosynthesis, which should balance the pro-thrombotic activity of blood platelets reported in MS (Saluk-Bijak et al., 2019). If so, genetic polymorphisms that reduce the enzymatic capacity or stability of this protein may constitute a genetic risk factor for MS, and serum may provide an ideal matrix to probe this variant association due to the activation of the 12-LOX pathway. 12/15 LOX showed the sharpest difference between control and MS cases which contains 14-HdoHE, 12-HEPE, and 12-HETE (**Figs. 4 & 8**). Of note, RRMS patients had higher levels of the pro-inflammatory mediator (15d-PGJ2, PGE3, and LTB5) and lower levels of 14-HDoHE, a pathway marker of maresin biosynthesis, a pro-resolving lipid mediator (Psychogios et al., 2011) and EPA-derived 12-HEPE, suggestive of lipid mediator imbalance in MS (**Figs. 2, 3, 7, & 8**). Adjustment of the values of a 12-LOX metabolite may alter the effect. This perhaps indicates the reduced activity of 12-LOX in the patients that impacts maresin production, which is in line with a previous study that found 12-LOX activity in platelets to be responsible for driving the precursor (14-HDoHE) formation leading to MaR1 production (Abdulnour et al., 2014). While ALOX12 variants are known, to the best of our knowledge they have not been explored in the context of this neuroinflammatory disease (Zheng et al., 2020).

Beyond 12-LOX metabolism, soluble epoxide hydrolase-dependent 12,13-DiHOME and its alpha-linolenic acid analog 12,13-DiHODE were both positively related to higher EDSS scores independent of age. Linoleate-derived 12,13-DiHOME, 9,10-DiHOME, and 9,12,13-TriHOME can be produced by 15-LOX in eosinophils (Fuchs et al., 2020). Little is known about the TriHOMEs, which are reportedly autoxidative products that can be reduced *in vivo* by dietary antioxidants (Picklo & Newman, 2015). The elevated presence of these metabolites may therefore indicate elevated oxidative stress in MS patients (Fuchs et al., 2018). Additionally, a reduced level of 10,11-EpDoPE was associated significantly with higher disease duration. Thus, the findings from our study indicate that the serum lipid mediator profiles, including some involved in SPM biosynthesis, represent specific features of inflammation and its resolution in MS. Overall, our analysis shows a distinct imbalance of pro-inflammatory and pro-resolving lipid mediators in RRMS patients compared to HS.

The balance between pro-resolution and pro-inflammatory mediators plays a crucial role in immune-mediated responses in inflammatory disease models (Serhan, 2014). Several studies have reported altered levels of PUFA metabolites in MS and EAE models (Derada Troletti et al., 2021; Kooij et al., 2020; Mangalam et al., 2013; Poisson et al., 2015; Pruss et al., 2013; Sanchez-Fernandez et al., 2022; Zahoor et al., 2023). The current findings also suggest impairment in the resolution pathway which may suggest altered functions exerted by these metabolites in governing the inflammatory process and its timely resolution. Previously, our group was the first to test the therapeutic potential of SPM (resolvin-RvD1) on disease progression in a preclinical animal model of MS-EAE (Poisson et al., 2015). Similarly, we have shown the protective action of maresin 1 in EAE by reducing inflammation, promoting neuroprotection, and preventing disease progression (Zahoor et al., 2023). Daily oral and intraperitoneal administration of RvD1 (100 ng per mouse) decreased disease progression to a greater extent and attenuated clinical symptoms by suppressing both peripheral and central inflammation. This study provided the first experimental proof that a small endogenous lipid mediator can act as therapy with drug-like properties in a mouse model of MS. Consistent with this work, a recent report confirmed the protective actions of DHA metabolites (resolvin-RvD1 and protectin-PD1) on inflammation-induced blood-brain barrier (BBB) dysfunction in MS using the human brain endothelial cell line hCMEC/D3 as a model system (Kooij et al., 2020). Treatment with these metabolites attenuated inflammation-induced BBB dysfunction, and activation as well as transendothelial migration of monocytes. Collectively, these studies provide crucial evidence supporting the therapeutic potential of DHA-metabolites in MS.

Since we found higher 15d-PGJ2 levels in RRMS patients, there is a study that reported *in vivo* treatment with 15d-PGJ2, a peroxisome proliferator-activated receptor-gamma (PPARγ) agonist and downstream lipid mediator of AA and PGD2-synthase, ameliorated EAE by blocking IL-12 signaling through the Janus kinases (JAK) signal transducer and activator of transcription (STAT) pathway and promoting Th1 differentiation (Natarajan & Bright, 2002). A second report demonstrated that prostaglandin synthesis through the mPGES-1-PGE2-EPs axis of AA cascade may exacerbate the disease pathology in EAE (Kihara et al., 2009), justifying higher levels of PGs in our study. To the best of our knowledge, there are only a few studies on SPMs in MS based on serum/serum and cerebrospinal fluid of study subjects (Kooij et al., 2020; Pruss et al., 2013) and the lipidomics data indicates diverse outcomes in the metabolite profiles in patient cohorts across studies, likely reflecting the impact of other physiological, dietary, and genetic factors and their interactions as modulators of metabolite levels. One such study is by Kooij *et al*., where authors could not detect maresin in serum samples (Kooij et al., 2020), however, they reported higher levels of its pathway marker 14-HDoHE, which contrasts with our study where we found lower levels of 14-HDoHE in patient samples. Also, we noted that ALA had higher intensity in RRMS samples compared to HS and may reflect an anti-inflammatory diet pattern in the RRMS vs HS populations (Gabbs et al., 2015; Takic et al., 2022). These sets of data provide the biological relevance of the altered lipid mediator metabolism in MS compared to HS, showing a direct correlation with heightened inflammation due to an imbalance between pro-inflammatory and pro-resolving mediators.

In conclusion, the findings from our study provide comparative profiling of lipid mediator signatures in serum samples of MS patients and HS. In agreement with the pre-clinical literature (Poisson et al., 2015; Zahoor et al., 2023), our study provides further evidence favoring an imbalance between the inflammatory response and resolution process in MS, which could be responsible for the partial recovery of the patients. Accordingly, there are two possibilities explaining this imbalance; either a defect in the synthesis of pro-resolution lipid mediators/metabolites or the rate of degradation of metabolites is higher than controls. Also, data quality controls indicate that analytical variability did not differentially influence metabolite levels in these samples, notwithstanding changes associated with sample collection or storage (La Frano et al., 2018). Consequently, it would appear that identifying how resolution mechanism(s) become disrupted will provide a better understanding of disease progression and aid in designing new therapies that can slow down inflammatory activation hasten the resolution of inflammation, and simultaneously halt disease progression. At the same time, our study has some limitations as we did not have information on parameters such as body weight/height, body mass index (BMI), adiposity, and lipemia; which could be possible confounders affecting the levels of omega fatty acids in blood. Further prospective longitudinal studies covering a large sample size are needed for generalized oxylipin and SPM profiling to clarify the levels of pathway markers for lipid mediator biosynthesis and their correlation with disease severity and disability score EDSS, thereby influencing disease course. Also, the effect of an altered key marker of maresin synthesis, 14-HDoHE on MS preclinical mouse models needs to be further validated. Overall, our study shows a strong relevance of decreased inflammation resolving lipid mediators and SPMS in RRMS patients and indicates that a strategy for developing SPMs as potential therapeutics may be a better approach compared to PUFA supplementation.

## Author Contributions

IZ compiled the manuscript; ID performed bioinformatics analysis; TP performed quantitative analysis, and data quality assessment and participated in biostatistics analysis, JW performed bioinformatics analysis and machine learning of data, MC, NA, LP, SMP, RR, and AYT, contributed to the design and finalized the manuscript; JWN and NA guided bioinformatics analysis, organize and finalized the manuscript; SG conceived the idea, directed the study, designed experiments, and reviewed the manuscript before approval for submission.

## Compliance with Ethical Standards Conflict of Interest

The authors declare that they have no conflict of interest.

## Supplementary data

**Supp. Fig 1:**
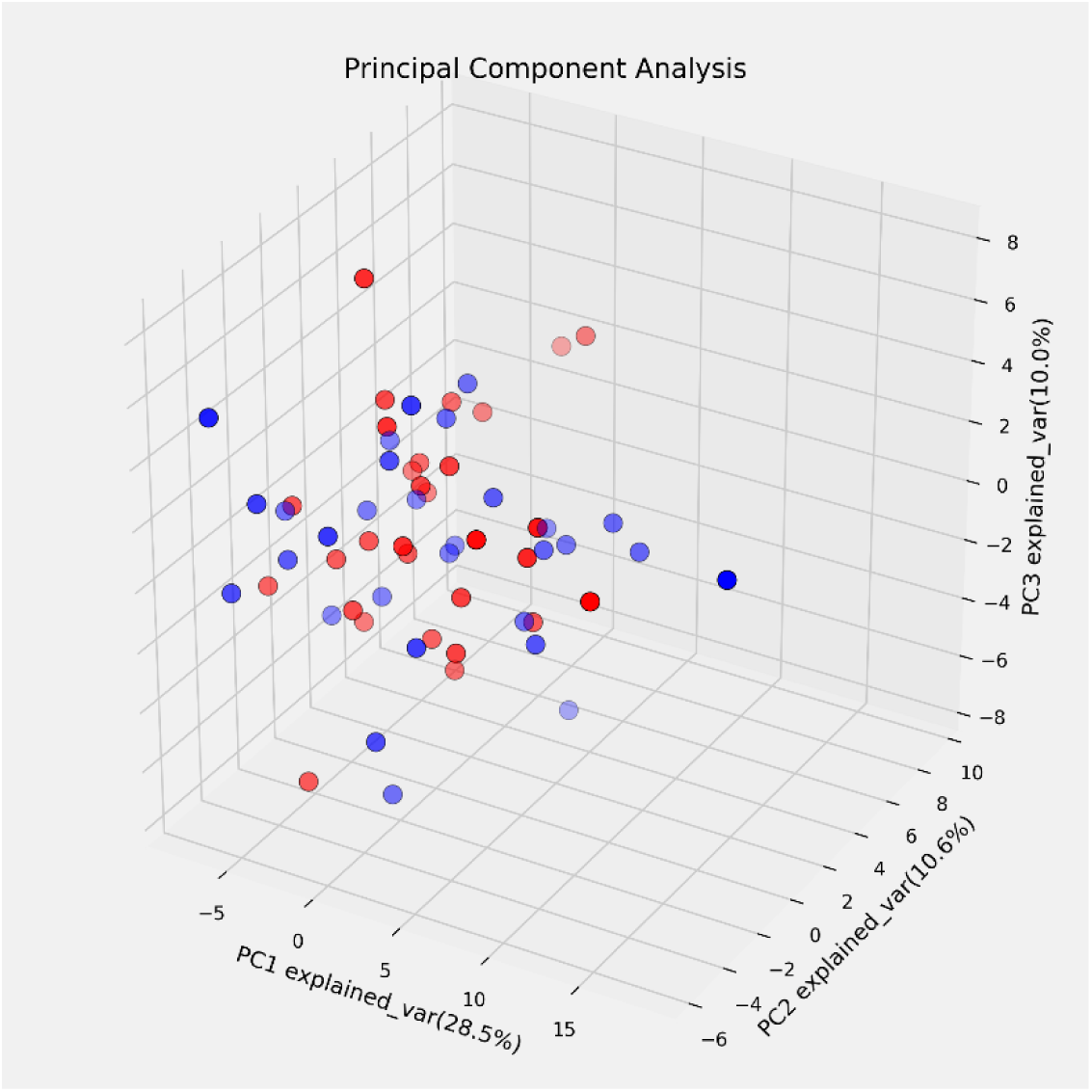
PCA Score plot of HS and RRMS samples. HS is labeled blue, RRMS red.

**Supp. Fig 2:**
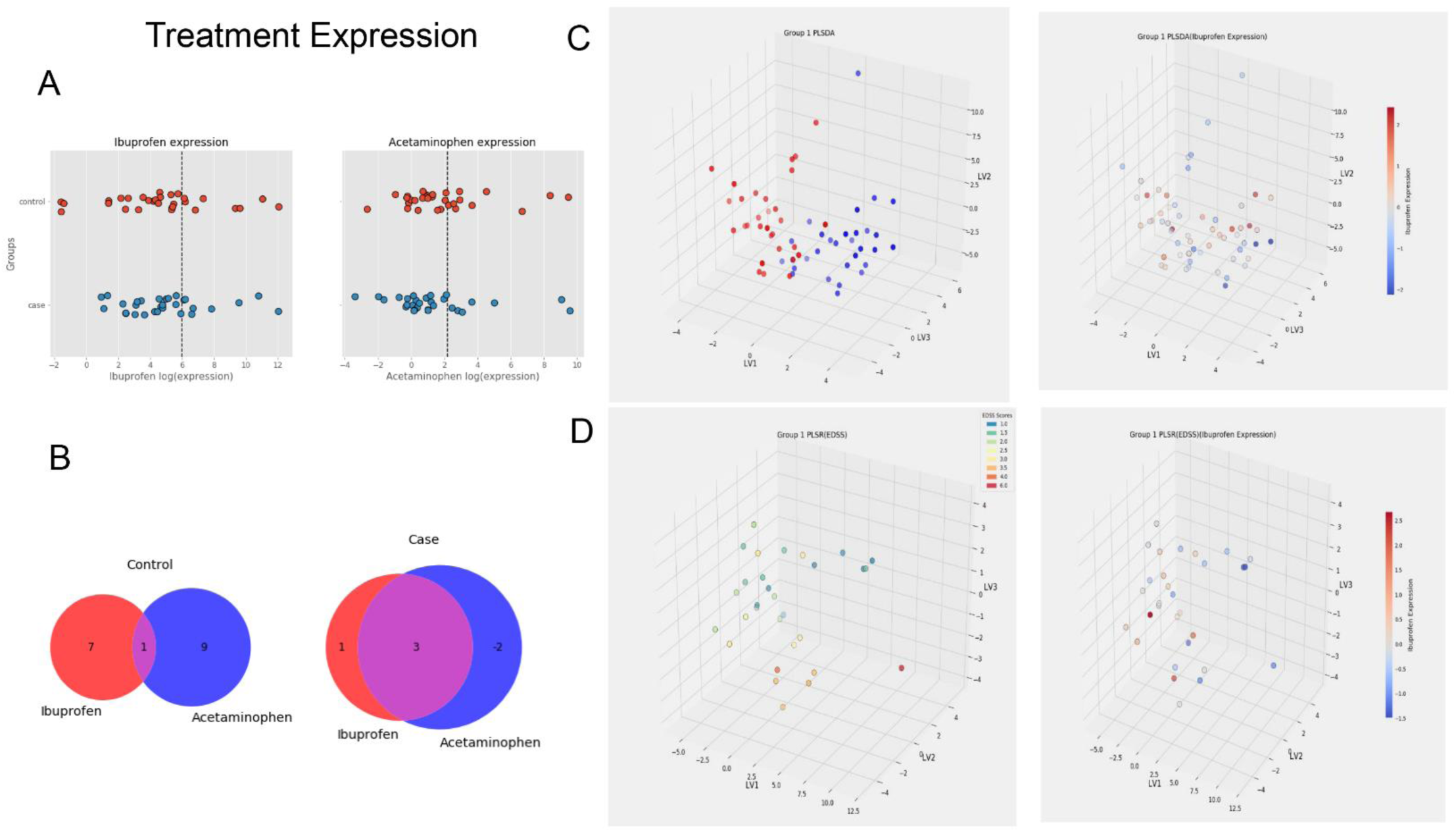
Show the values of Ibuprofen and acetaminophen (log(nM)) in the plasma of HS and RRMS subjects. The dashed line in **2A.** Corresponds to high levels of Ibuprofen (6 nM) and acetaminophen (2.2 nM). **2B.** Venn diagram displaying several subjects with high levels of Ibuprofen/Acetaminophen. **2C** and **2D** PLSDA and PLSR, respectively, showed samples by the group for PLSDA (HS: blue, RRMS: red) and EDSS score for RRMS samples for PLSR. Juxtaposed are score plots labeled by ibuprofen expression.

